# Varying phylogenetic signal in susceptibility to four bacterial pathogens across species of Drosophilidae

**DOI:** 10.1101/2024.04.19.590331

**Authors:** Hongbo Sun, Mark A. Hanson, Sarah K. Walsh, Ryan M. Imrie, Ben Raymond, Ben Longdon

## Abstract

Bacterial infections are one of the major sources threatening public health. Pathogen host shifts – where a pathogen jumps from one host species to another – are important sources of emerging infectious diseases. However, compared to viruses, we know relatively little about the factors that determine whether a bacteria can infect a novel host, such as how host phylogenetics constrains variation in pathogen-host range and the link between host phylogeny and the infectivity and virulence of a pathogen. Here, we experimentally examined susceptibility to bacterial infections using a panel of 36 Drosophilidae species and four pathogens (*Providencia rettgeri*, *Pseudomonas entomophila*, *Enterococcus faecalis*, *Staphylococcus aureus*). The outcomes of infection differed greatly among pathogens and across host species. The host phylogeny explains a considerable amount of variation in susceptibility, with the greatest phylogenetic signal for *P. rettgeri* infection, explaining 94% of the variation in mortality. Positive correlations were observed between mortality and bacterial load for three out of the four pathogens. Correlations in susceptibility between the four pathogens were positive but largely non-significant, suggesting susceptibility is mostly pathogen-specific. These results suggest that susceptibility to bacterial pathogens may be predicted by the host phylogeny, but the effect may vary in magnitude between different bacteria.

## Introduction

Emerging and re-emerging infectious diseases are a major threat to public health [1–3]. Multi drug-resistant bacterial infections are a major cause of mortality, responsible for millions of deaths globally [4]. Recent studies have highlighted that host shifts – where a pathogen jumps from one host species to another – are important sources of emerging diseases, not only by viruses, but also by bacterial pathogens [5, 6]. There is an increasing awareness that bacterial pathogens also commonly undergo host shifts [6–8], which has been linked to their recent emergence as causative agents of disease in several host systems, including humans, livestock, wildlife, and plants [7–9]. In *Staphylococcus aureus*, a human pathogen associated with several severe hospital and community-acquired infections [10], humans are the primary transmission hub for host shifts into a range of livestock species, and cows have been considered the main animal reservoirs for human epidemics [11–13]. Another extensively studied example is the jump of *Mycoplasma gallisepticum* from poultry to wild house finches [14]. Here, the *Mycoplasma* host shift has driven the evolution of resistance in the house finch hosts [15], led to changes in bacterial gene content [7] and the evolution of virulence [16]. In plants the bacterial pathogen *Xylella fastidiosa* is found in over 600 host species [17]. A single gene loss event is associated with the host shift of *X. fastidiosa* from coffee plants to olive trees [18]. Most studies on bacterial host shifts are based on genomic data following natural host shifts, and there are limited experimental studies of bacterial host shifts that move beyond two-host systems.

Perhaps the best experimental studies relating to bacterial host shifts come from insects and their bacterial endosymbionts. In a study of coccinellid beetles and a male-killing isolate of *Spiroplasma*, the bacterium’s ability to distort offspring sex ratios in novel hosts was negatively correlated with the genetic distance from the bacteria’s natural host [19]. Similar patterns have been found for *Wolbachia*, a bacterial endosymbiont that infects ∼40-60% of all arthropods [20–22]. Artificial transfers of *Wolbachia* are most successful between closely related hosts [23, 24]. This tallies with field data on *Wolbachia* distributions that found strong effects of the host phylogeny, with high rates of *Wolbachia* sharing between closely related species, and a decline in co-occurrence in different hosts with increasing phylogenetic distance between these hosts [9]. Viruses show similar trends, with phylogenetic distance and clade effects explaining large amounts of variation in susceptibility to several viruses [25, 26]. When comparing different viruses from the same family, positive inter-specific correlations in susceptibility have been observed, suggesting similar mechanisms underlie differences in susceptibility observed among species [26, 27]. Whether such patterns hold true for free-living bacteria, which are less tightly linked to their hosts, remains to be tested.

Here, we have used a large cross-infection experiment to understand how susceptibility to bacterial pathogens varies across host species. We infected a large panel of 36 species of Drosophilidae with four bacterial pathogens: two Gram-positives, *Staphylococcus aureus*, *Enterococcus faecalis*, and two Gram-negatives: *Providencia rettgeri* and *Pseudomonas entomophila*. In *Drosophila melanogaster*, the major immune response to bacterial pathogens involves the Toll and IMD pathways. The Toll pathway primarily controls resistance to fungal and Gram-positive bacterial infections, whereas the Imd pathway mediates response to Gram-negative bacterial infections [28, 29]. Both of these *NF-κB* pathways result in the upregulation of antimicrobial peptides (AMPs) and other defence response genes [30, 31]. Genes in these pathways evolve rapidly. However, it remains unclear how differences in immune genes among species contributes to differences in infection outcome [32, 33]. Recent studies have demonstrated that AMPs show specific importance for certain bacterial infections and that different alleles of AMPs are linked to differences in susceptibility both within and across species [34, 35].

The bacterial pathogens used here have been well characterised in *D. melanogaster* [36, 37]. *E. faecalis* was initially derived from a human clinical isolate [38, 39]. The virulence of *E. faecalis* varies with different strains [40]. The *E. faecalis* FA2-2(pAM714) used in this study produces toxins (e.g. cytolysin), that act as virulence factors to improve bacterial colonisation and decrease host survival. in *Drosophila* [39], the Toll pathway, particularly the host defence peptides Bomanin and Baramicin, have been shown to play a role in response to *E. faecalis* [41–43]. *S. aureus* was initially isolated from humans and has been used in mice and *Drosophila* infection models [36, 44]. *S. aureus* is a multispecies pathogen that can cause disease in humans and livestock. The dynamic gain and loss of host-specific adaptive genes, usually located on mobile genetic elements (MGEs), enable *S. aureus* to jump between different species [13, 45]. In *D. melanogaster*, resistance to *S. aureus* specifically relies on the melanisation response [46, 47]. *P. entomophila* was isolated from a wild female *D. melanogaster* and is a versatile soil bacterium that can infect *D. melanogaster* as well as insects from different orders. Upon infection, *D. melanogaster* show differential expression of genes including AMPs, oxidative stress response genes and detoxification genes [48–51]. Genes involved in host damage prevention, signalling, renewal, and regulation also affect the susceptibility of *D. melanogaster* when *D. melanogaster* is orally challenged with *P. entomophila* [52]. *P. entomophila* causes digestive obstruction and eventually leads to the host’s death by damage and perforation of the gut through the production of insecticidal protease-based toxins [49, 51]. *P. rettgeri* was initially isolated from the haemolymph of wild *D. melanogaster* [53]. The IMD pathway, and particularly the AMP Diptericin has been shown to be an important part of the immune response to *P. rettgeri* [35, 54, 55]. A single nucleotide polymorphism at residue 69 in *Diptericin A*, “S69R” of *DptA*, is associated with increased susceptibility to *P. rettgeri* bacterial infection [34, 35].

Here, we examine the role of the host phylogeny in determining the outcomes of bacterial infections. We investigated how variation in bacterial virulence varies across the host phylogeny as well as the relationship between virulence and bacterial load for each of the four bacterial pathogens.

## Methodology

### Bacterial strains and culture

We used four bacterial pathogens in this study. Two Gram-positive bacteria, *Staphylococcus aureus* PIG1 (*S. aureus*) [44] and *Enterococcus faecalis* FA2-2/pAM714 (*E. faeclis*) [38, 39], and two Gram-negative bacteria: *Providencia rettgeri* Dmel (*P. rettgeri*) [53] and *Pseudomonas entomophila* (*P. entomophila*) [49]. Bacterial cultures were initiated from frozen glycerol stocks onto LB (Lysogeny Broth) agar, with the exception of *P. entomophila,* which were plated on egg yolk agar (10g of agar, 10g of LB, 1 L of water and 20 mL of egg yolk (Sigma)). Colonies of *P. entomophila* with halos on egg yolk agar (indicating phospholipase activity) were used to prepare inocula. Insect inocula were cultured in LB broth, with the exception of *E. faecalis* which used BHI (Brain heart infusion) broth. All bacteria were grown at a 30℃ incubator, shaking 180 rpm/min for liquid culture. Inocula were prepared in a three-step process in order to standardise physiology in the stationary phase. Initially, colonies were used to prepare cultures of 5 mL in 30 mL sterile universals. These were diluted in liquid media to OD_600_ 0.2, and then 50 µL of this was inoculated into new vials. Following overnight growth, cultures were diluted with culture media to the desired OD value (OD 2.0 for *E. faecalis* and OD 1.0 for the other bacteria based on pilot mortality data).

### Fly stocks and rearing

All flies were maintained in multi-generation stock bottles (Fisher brand) at 22°C, ∼50% relative humidity in a 12-hour light-dark cycle. Each stock bottle contained 50 ml of one of four varieties of *Drosophila* media which were chosen to optimise rearing conditions (full details in Supplementary Table 1). Male flies were used to avoid any effect of sex or mating status, which has been shown to influence the susceptibility of female flies to various pathogens [56–59]. Before inoculation, 0-1 day old male flies were transferred to vials containing cornmeal media. These flies were then transferred to fresh media every three or four days for a week (age 7–8 days, flip once before infection), at which point they were inoculated. Flies were kept at 22°C, 70% relative humidity in a 12-hour light-dark cycle throughout the experiments. Flies were inoculated under CO_2_ anaesthesia via septic pin prick with 10μm diameter stainless steel needles (<u>26002-10</u>, Fine Science Tools, CA, USA). These needles were bent approximately 250μm from the end to provide a depth stop and dipped in the inoculum before being pricked into the anepisternal cleft in the thorax of anaesthetised flies. Inoculation by this method bypasses the gut immune barrier but avoids differences in inoculation dose due to variations in feeding rate in oral inoculation. This method has been demonstrated to be highly repeatable (see results) and gives a consistent initial dose (our estimate of initial dose (CFUs) introduced into individual fly for each pathogen: OD = 1 *P. rettgeri* :10^(1.94±0.08); OD = 1 *P. entomophila* : 10^(1.50±0.14); OD= 2 *E. faecalis* :10^(2.67±0.12); OD = 1 *S. aureus* : 10^(2.75±0.12). All inoculations were done by a single experimenter (HS).

For the survival assay, we inoculated male flies of 36 fly species. We inoculated three replicate vials for 165 species-bacteria combinations, two replicate vials for 13 combinations and one replicate vial for two combinations. For each combination, one of 36 species was infected with either a control Ringer’s solution [60] or one of the four bacterial pathogens listed above. Vials contained a mean of 14 flies (range = 5-18 flies). Following pin prick infection, the number of dead flies in each vial was recorded daily for 14 days. Flies were tipped into new vials of food every two days.

We measured bacterial load in 18 species of fly; these species were selected based on their survival results and covered a broad range of survival values for each pathogen. We measured colony forming units (CFUs) for individual male flies of each species (mean = 15 replicate flies, range 1-16) infected with each species-bacteria combination (there is only one replicate for *D. teisseri* in *P. entomophila* infection (Figure 3) as most of flies died due to infection before collection, there are at least 7 flies per species with a mean of 15 flies for all other treatments.). CFUs were measured at 16 hours post-infection. Flies were prepared and infected as above. Before homogenisation, anaesthetised flies were moved to 96-well tubes containing a 4mm steel ball and 200 µL 70% ethanol, flies were then gently cleaned using 70% ethanol and homogenised in 200 µL sterile LB or sterile BHI (for flies infected with *E. faecalis*) in a Tissue Lyser II at 30 Hz for three minutes. Homogenates were diluted 10^-1^ to 10^-5^ and plated on LB agar or BHI agar in two technical replicates before being cultured overnight at 30℃. Bacterial loads were measured as colony-forming units (CFUs) per fly, following the formula: CFU per fly = (Colony counts × Dilution factor × Homogenate volume) / Plating volume.

### Host Phylogeny

The method used to infer the host phylogeny has been described elsewhere [61] and details are provided in the supplementary methods.

### Sequencing of *Diptericin A*

To determine the *Diptericin A* allele of each species (which has been linked to susceptibility to bacterial pathogens [35]), *Diptericin* genes were sequenced across species to determine their S69R allele (see supplementary methods).

### Statistical analysis

We used phylogenetic mixed models to look at the effects of host species relatedness on bacterial susceptibility across the 36 host species. We fitted all models using a Bayesian approach in the R package MCMCglmm v2.29 [62] in RStudio (R v4.2.2) [63]. We used a multivariate model with mortality following inoculation with each of the four bacteria or control treatments as the response variables. Mortality was calculated as each vial’s mean proportion of dead flies over 14 days.

The models took the following form:

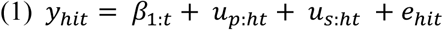

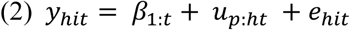

Where 𝑦 is the mortality of the *i^th^* biological replicate of host species *h,* for treatment *t* (each of the four bacteria and the non-infected control). *β* are the fixed effects, with *β_1_* being the intercepts for each trait.

We also ran bivariate models for each of the four pathogens across the 18 host species tested to investigate how mortality was related to bacterial load. These models had the following form:

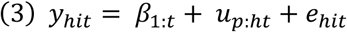

Where 𝑦 is the response of the *i^th^*biological replicate of host species *h,* for treatment *t* (either mortality or the bacterial load (CFUs) of an individual fly infected with that pathogen). *U* are random effects for the phylogenetic species effects (*p*), non-phylogenetic species effects (*s*) and residuals (*e*). 𝑢_𝑠:ℎ𝑡_was removed from models (2) and (3) as the models struggled to separate the phylogenetic and non-phylogenetic effects. The random effects and the residuals are assumed to be multivariate normal with a zero mean and a covariance structure ***V_p_*** ⊗ **A** for the phylogenetic effects and ***V_e_***⊗ **I** for the residuals (⊗ here is the Kronecker product). **A** is the phylogenetic relatedness matrix, **I** is an identity matrix and the **V** are 5×5 (co)variance matrices describing the (co)variances between mortality when infected with each of the four bacteria or the control. In model (3) the **V** are 2×2 (co)variance matrices describing the (co)variances between mortality and bacterial load when infected with each of the four bacteria. Specifically, the matrices ***V_p_*** and ***V_s_*** describe the phylogenetic and nonphylogenetic between-species variances in mortality for each treatment and the covariances between them, whereas the residual covariance matrix ***V_e_*** describes within-species variance that includes both actual within-species effects and measurement error. The off-diagonal elements in ***V_e_*** (the covariances) are unable to be estimated because no vial has multiple measurements – so they were set to zero. Model (1) was also run with two additional fixed effects of body size and *Diptericin A* S69R site status (see supplementary methods).

The MCMC chain was run for 13 million iterations with a burn-in of 3 million iterations and a thinning interval of 5000 iterations. We confirmed our experimental design and sample sizes had sufficient power to detect effects by down sampling a similar dataset [27]. The results presented were obtained using parameter expanded priors for the **V**_p_ and **V**_s_ matrices [66]. Results were tested for sensitivity to the use of different priors by being run with different prior structures. The alternative priors gave qualitatively similar results, except for the correlations where we found mean estimates were higher with the majority being significant. To determine what may be driving this we simulated response data based on mean estimates from the parameter expanded prior model using the ‘simulate.MCMCglmm’ function. We then refitted models on this data with either parameter expanded or alternative priors (inverse wishart or flat). The alternative priors had higher estimates of the correlations in the refitted models suggesting they may be inadvertently influencing the estimates. The proportion of the between-species variance that the phylogeny can explain was calculated from model (1) using the equation ***V_p_ /(V_p_ + V_s_)***, where ***V_p_*** and ***V_s_***represent the phylogenetic and species-specific components of between-species variance [67], respectively, and are equivalent to phylogenetic heritability or Pagel’s lambda [68, 69]. The repeatability of bacterial load measurements was calculated from model (2) as ***V_p_/(V_p_ + V_e_)***, where ***V_e_*** is the residual variance of the model [70]. Interspecific correlations were calculated from model (2) and (3) ***V_p_*** matrix as ***cov_x_,_y_ / √(var_x_ × var_y_)***. Parameter estimates reported are means of the posterior density, and 95% credible intervals (CIs) were taken to be the 95% highest posterior density intervals.

We ran additional models using Abbots correction on data, and a binomial model on day 14 mortality (see supplementary methods). We confirmed our results were robust to these different analyses (supplementary Table S5-S6, Figure S3) and so results presented are the main analysis described above.

## Results

### Patterns of susceptibility to bacterial pathogens across host species

To investigate how susceptibility to bacterial pathogens varies across host species, we measured survival in 7402 flies from 36 species of Drosophilidae inoculated with one of four bacterial pathogens or a control. We found large differences in the mean virulence of the bacteria; the mean mortality was 70% for *P. entomophila* infection, 46% for *P. rettgeri*, 39% for *S. aureus*, and 36% for *E. faecalis* (compared to only 4% in control inoculated flies). There were dramatic differences in the survival of the different species with the majority of individuals of certain species dying, whilst others showed little mortality when infected with the same pathogen (Figure 1). For example, just over half of the species showed rapid mortality following *P. entomophila* infection, with over 50% of flies dying within one day post-infection in 19 out of 36 fly species (Figure 1). Conversely, survival remained high for most species infected with *E. faecalis,* with the exceptions of *D. yakuba* and *D. sucinea* where >85% of flies had died by the third day post-infection (Figure 1).

**Figure 1.**
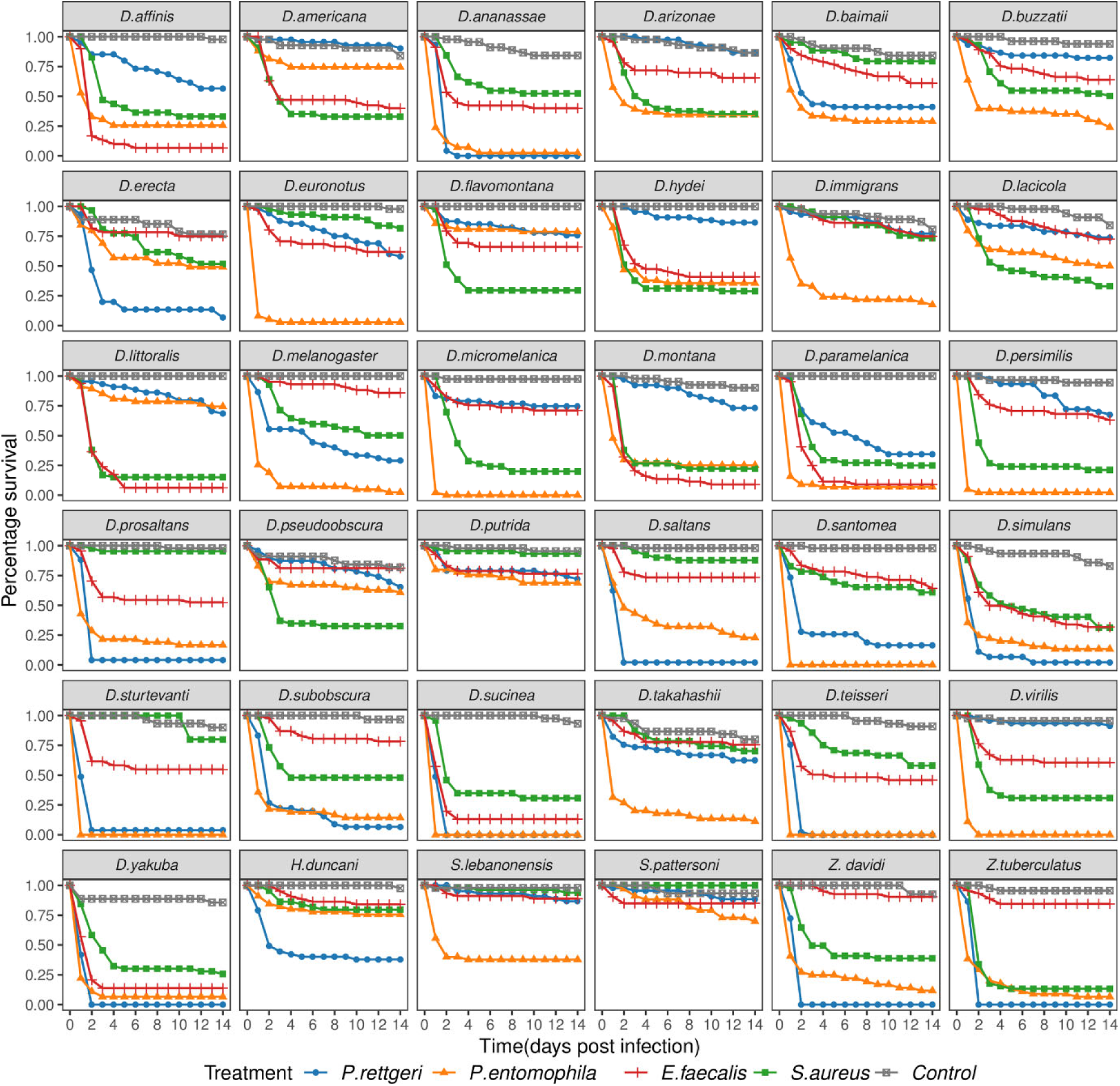
Survival of 36 species of Drosophilidae after infection with four bacterial pathogens. Each point represents the mean survival of three replicates, colours represent the inoculation treatment.

### The host phylogeny explains varying amounts of variation in mortality

Different clades of hosts show different mortality levels with each bacteria, with hosts with high and low mortality clustering together on the host phylogeny (Figure 2). For example, a clade within the subgenus Drosophila (delineated by *D. micromelanica* down to *D. flavomontana* in Figure 2) shows low mortality when infected with *P. rettgeri* compared to the other major host clades. We used a phylogenetic mixed model to determine the proportion of variation in mortality explained by the host phylogeny for each pathogen (Table 1). The estimates vary among the four pathogens, with the host phylogeny explaining large amounts of variation for three out of four bacterial infections. We find that almost all of the variation in mortality is explained by the phylogeny following *P. rettgeri* infections (94%).

**Figure 2:**
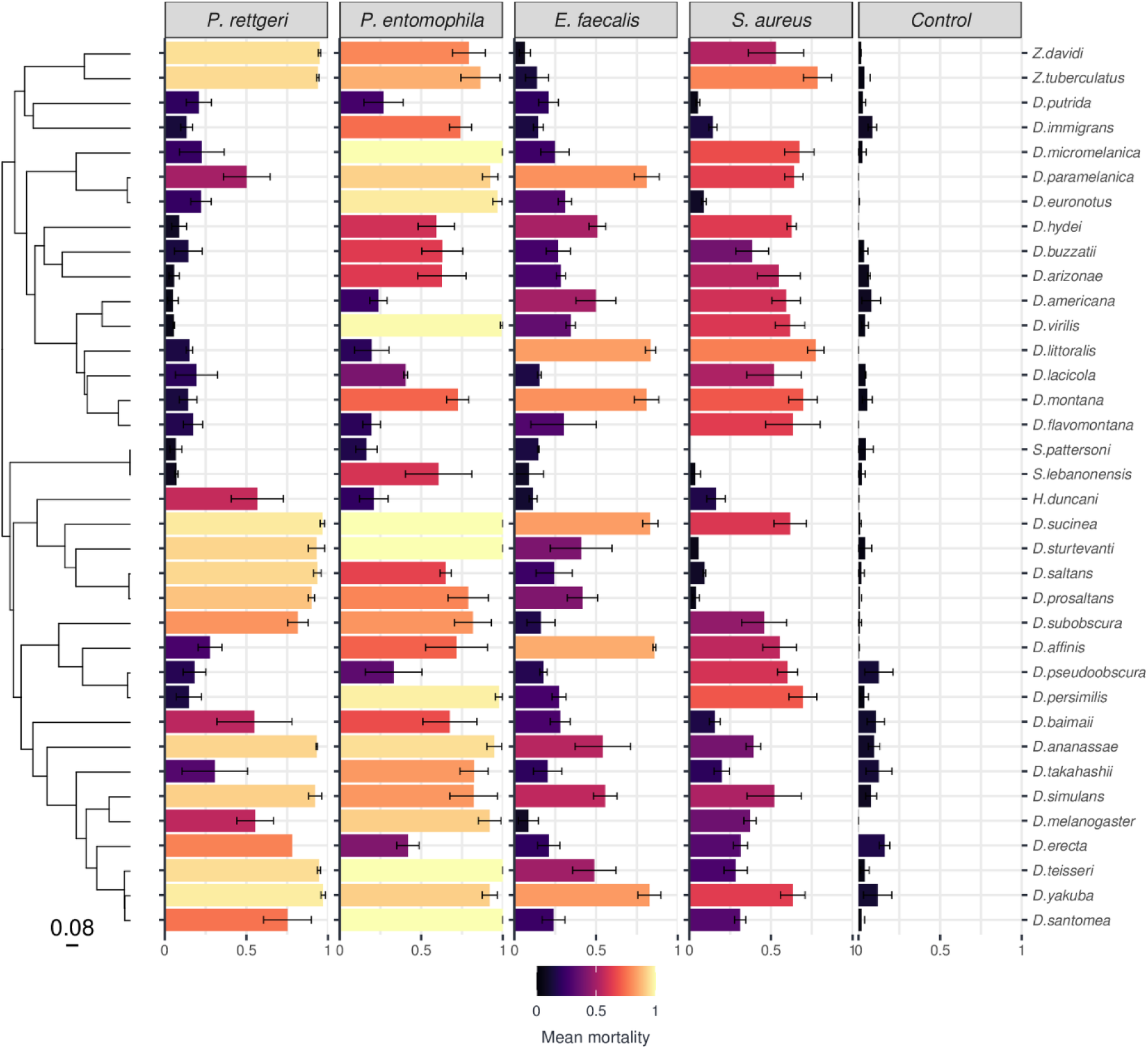
Susceptibility to four bacterial pathogens varies across 36 Drosophilidae species. Bar length and colour represent mean mortality change across pathogen and fly species with error bars representing standard errors. The phylogeny of Drosophilidae host species is presented on the left, with the scale bar representing the number of nucleotide substitutions per site. Names of host species are given on the right; the column label shows different treatments (bacterial or control inoculation).

**Table 1.** Estimates of the mean proportion of variation explained by the host phylogeny (taken from model (1)), and the repeatability of estimates (taken from model (2)).

| Pathogen | Variance explained by phylogeny | Repeatability |
| --- | --- | --- |
| *P. rettgeri* | 0.94 (95% CI: 0.81, 1.00) | 0.84 (95% CI: 0.75,0.93) |
| *P. entomophila* | 0.36 (95% CI: 0.00, 0.74) | 0.70 (95% CI: 0.48,0.89) |
| *E. faecalis* | 0.14 (95% CI: 0.00, 0.50) | 0.90 (95% CI: 0.80,0.97) |
| *S. aureus* | 0.64 (95% CI: 0.24, 0.94) | 0.80 (95% CI: 0.63,0.92) |

The phylogeny explains a substantial proportion of the variation in mortality for *S. aureus* (64%) but less for *P. entomophila* (36%) and *E. faecalis* (14%). However, these have broad 95% CI’s around estimates of the posterior mean, which are pushed up against zero for *P. entomophila* and *E. faecalis*. If we compare the phylogenetic signal in mortality for each of the pathogens we find that the phylogeny explains a greater amount of variation in mortality to *P. rettgeri* infection than for the other pathogens (although the effect is marginal for *P. rettgeri - S. aureus*), and *S. aureus* has greater phylogenetic signal than *E. faecalis* (Table S2). Measurements were highly repeatable (Table 1) with a large proportion of the variance being explained by the inter-specific phylogenetic component, with little within species variation or measurement error (Repeatability = *V_p_/(V_p_ + V_e_)*).

### Patterns of susceptibility are largely distinct for each bacteria

In *D. melanogaster*, the host immune response differs markedly between infections by Gram-positive and Gram-negative bacteria [71, 72]. If mechanisms of host defence depend on the induced immune response, we might expect positive correlations in host susceptibility among species for bacteria of the same Gram type. We found the correlations between bacteria all had positive estimates (Table 2), although the 95% CIs overlapped zero for all except for the correlations between *S. aureus–E. faecalis* and *E. faecalis–P. rettgeri.* The correlation coefficient was less than 0.4 for all correlations bar the estimate between the two Gram-positive bacteria *S. aureus* and *E. faecalis* where *r* = 0.66. Overall, this suggests that although hosts may show similar trends in susceptibility to all the bacteria tested (correlations are largely positive); host susceptibility generally differs by pathogen, with shifts in the rank order of species susceptibilities to the different pathogens (i.e. as correlations are all less than one, so species do not show overall high or low susceptibity to bacteria *per se*, but rather pathogen specific susceptibilities).

**Table 2.** Correlations in mortality between different pathogen and control treatments. Numbers represent the mean estimates of correlation coefficients from Model (2) with 95% credible intervals shown in brackets. Significant correlation coefficients (with 95% CI that do not span zero) are highlighted in bold.

| Bacteria | Inter-specific correlation $r$ |
| --- | --- |
| *S. aureus* - *E. faecalis* | **0.66 (95% CI: 0.37, 0.90)** |
| *S. aureus* - *P. rettgeri* | 0.35 (95% CI: -0.04, 0.70) |
| *S. aureus* - *P. entomophila* | 0.34 (95% CI: -0.13, 0.73) |
| *E. faecalis* - *P. rettgeri* | **0.39 (95% CI: 0.02, 0.72)** |
| *E. faecalis* - *P. entomophila* | 0.31 (95% CI: -0.19, 0.74) |
| *P. rettgeri* - *P. entomophila* | 0.37 (95% CI: -0.05, 0.75) |
| *P. rettgeri* - Control | -0.06 (95% CI: -0.65, 0.61) |
| *P. entomophila* – Control | -0.04 (95% CI: -0.65, 0.66) |
| *E. faecalis* - Control | -0.15 (95% CI: -0.85, 0.66) |
| *S. aureus* - Control | -0.12 (95% CI: -0.80, 0.64) |

### Mortality correlates with bacterial load in most cases

To test whether increased mortality is due to higher bacterial loads, we selected 18 fly species that varied in their mortality and measured their bacterial load to each pathogen at 16 hours post-infection. We found that 24% of the flies infected by *P. entomophila* had died by 16 hours post-infection (<2% of flies infected with each of the other bacteria died by this time point). We excluded dead flies from the analysis below to avoid confounding bacterial growth rates, as these flies may have died prematurely from injury, and bacterial growth rates may have developed differently in carcasses or flies failing to mount a full immune response. however, we note that the results were robust if we included the bacterial load measured in corpses. We quantified bacterial load in 1328 individual flies (Figure 3) and ran models combining this with the mortality data above for these 18 fly species.

**Figure 3:**
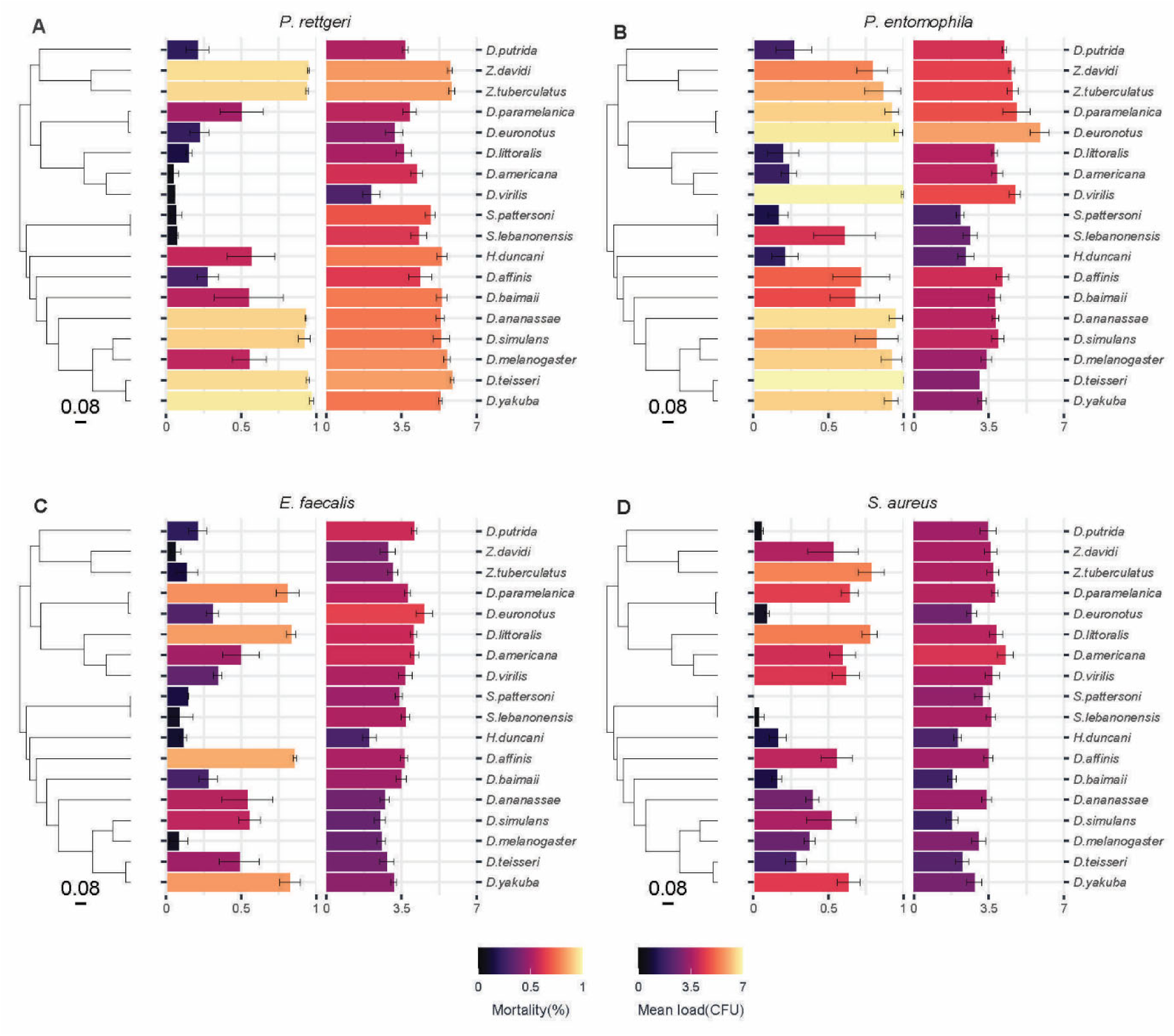
Mortality and bacterial load to four bacterial pathogens across 18 fly species. Bar length and colour represent mean mortality (left) or bacterial load (right) across 18 fly species bearing either *P. rettgeri* (A), *P. entomophila* (B), *E. faecalis* (C) or *S. aureus* (D) infection. Bacterial loads are data point at 16 hours post infection, with the error bars representing standard errors of a mean of 15 replicate flies for each species. The phylogeny of Drosophilidae host species is presented on the left, with the scale bar representing the number of nucleotide substitutions per site. Names of host species are presented on the right.

Mortality estimates from this model using a subset of the data from the survival experiment (Table 3) are consistent with observations in survival experiments (Figures 1 and 2). Mean bacterial loads are relatively consistent across the four pathogens (Table 3). We found strong positive correlations between mortality and bacterial load for *P. rettgeri* and *P. entomophila* and a slightly lower positive correlation for *S. aureus* with 95% CI’s that just overlap zero (Table 4, Figure S1). However, we found no evidence of a significant positive correlation between mortality and bacterial load for *E. faecalis* (Table 4, Figure S1). However, the 95% CI’s around estimates were broad so there was no clear indication that bacteria were behaving differently.

**Table 3.** Estimations of mortality and bacterial load for each pathogen. Numbers represent the mean estimates of mortality (mean proportion dead) and bacterial load (CFU per fly on a Log10 scale) for each pathogen from Model (3) with 95% credible intervals shown in brackets.

| Pathogen | Bacterial load | Mortality |
| --- | --- | --- |
| *P. rettgeri* | 4.63 (95% CI: 3.67, 5.50) | 0.46 (95% CI: 0.17, 0.76) |
| *P. entomophila* | 3.78 (95% CI: 2.97, 4.46) | 0.62 (95% CI: 0.33, 0.90) |
| *E. faecalis* | 3.36 (95% CI: 2.80, 3.92) | 0.37 (95% CI: -0.02, 0.74) |
| *S. aureus* | 3.12 (95% CI: 2.32, 3.85) | 0.36 (95% CI: 0.06, 0.68) |

**Table 4.** Correlations between mortality and bacterial load for each pathogen. Numbers represent the mean estimates of correlation coefficients from Model (3) with 95% credible intervals shown in brackets. Significant correlation coefficients are where the 95% CI do not span zero.

| Pathogen | Inter-specific correlation $r$ |
| --- | --- |
| *P. rettgeri* | 0.59 (95% CI's: 0.16, 0.94) |
| *P. entomophila* | 0.54 (95% CI's: 0.10, 0.90) |
| *E. faecalis* | 0.22 (95% CI's: -0.43, 0.74) |
| *S. aureus* | 0.48 (95% CI's: -0.03, 0.93) |

We found some evidence of microbiome proliferation in pathogen infected insects when measuring CFU in four species (*D. americana*, *S. lebanonensis*, *D. putrida* and *S. pattersoni*). We note there was an elevated count of likely microbiome-specific colonies in these species based on colony morphology and growth rate compared to colonies of pathogenic bacteria on LB/BHI agar. We included these colonies in our final bacterial load counts to avoid subjective calls on microbiome or pathogen colony identification, but will note that excluding these species from our analysis did not alter the results qualitatively.

### Testing for effect of *Diptericin A* allelic variation and body size

We observed an exceptionally high proportion of variance explained by host phylogeny for *P. rettgeri* (Table 1). This suggests some phylogenetically inherited factor, or factors, strongly determines susceptibility to this pathogen. Previous studies linked the gene *Diptericin A* (*DptA*) to defence against *P. rettgeri*, including resistant or susceptible alleles at a polymorphic residue both within and across species [35, 54].

We successfully amplified and sequenced *DptA* from 29 species and annotated the allele status at a previously identified polymorphic residue (residue 69 of *D. melanogaster DptA*). Four residues were found in our panel at this site (Figure S2). These were dominated by the resistant allele encoding serine (S) in 75% of species, and three alternate alleles each found in only 2-3 species, including a known susceptible allele encoding arginine (R), and alleles encoding either glutamine (Q) or limited replication of each individual allele, we fitted a fixed effect of *DptA* residue 69 by pathogen with species either being resistant (S and Q) or susceptible (R and N). We found no significant effect of the susceptible *DptA* alleles for any of the four pathogens (Table S4) but note there was limited power to detect an effect with only five species having susceptible alleles. There was little change in the variance explained by the phylogeny when the *DptA* allele status was included as a fixed effect (Table S4) indicating this allele does not readily explain the phylogenetic patterns observed.

The wing size of *Drosophila* can provide a proxy for body size [73] which is known to be phylogenetically dependent [74]. We included wing size as a fixed effect in models to see if wing size explained some of the variation in host susceptibility. We found no effect of wing size on susceptibility for any pathogen (mean and 95% CIs for each pathogen: *P. rettgeri*: −0.34 (95% CI:- 0.75, 0.02), *P. entomophila*: −0.02 (95% CI:-0.54, 0.45), *E. faecalis*: 0.01 (95% CI:-0.37, 0.37), *S. aureus*: 0.03 (−0.37, 0.42)).

## Discussion

We examined how susceptibility to bacterial pathogens varies across the host phylogeny, using a panel of 36 Drosophilidae species and four bacterial pathogens. Using phylogenetic mixed models we found the host phylogeny can explain a large proportion of variation in susceptibility, but differs in the extent of variation explained between pathogens. Distinct infection patterns were seen between the different pathogens, with all correlations in susceptibility between the pathogens being less than one, suggesting susceptibility generally varies between bacterial pathogens. However, despite being largely insignificant, all correlations had positive estimates, and the strongest correlation was seen between the two Gram-positive bacteria (Table 2) suggesting that differences in susceptibility may have shared underlying mechanisms. Positive correlations between mortality and bacterial load for *P. rettgeri*, *P. entomophila* and *S. aureus* suggest virulence is a direct result of bacterial accumulation in the host.

The lack of correlation between virulence and bacterial load for *E. faecalis* may have a biological underpinning, such as the production of toxins, or inhibited replication capabilities of the human *E. faecalis* strain we used, which may not be well adapted to invertebrate hosts [40]; alternately this may be due to limited statistical power. These results (Figure S1) also demonstrate that different species can have differences in mortality with similar bacterial loads. As such, examination of whether potential tolerance mechanisms underly differences in susceptibility [75] could be an interesting follow up to this work.

Our survival results find a substantial amount of variation in susceptibility can be explained by the host phylogeny, with the largest effect for *P. rettgeri,* where almost all of the variation in susceptibility is explained by the host phylogeny. One potential explanation is that susceptibility is linked to variation in immune genes with certain host clades having lost or gained different aspects of the immune system. Previous studies have shown that a specific AMP, *Diptericin*, plays an essential role in defence against *P. rettgeri* infection. Flies encoding arginine (R) at the *Diptericin* S69R polymorphic site are more susceptible than flies encoding serine (S) [35, 54]. Due to the limited statistical replication of the different *Diptericin* alleles (75% of the 29 species with allele data had the “S” resistance allele, with each of the other 3 alleles only replicated in 2 or 3 species) we had little power to detect effects here. However, there is also substantial variation in susceptibility just among species with the “S” allele, suggesting this *Diptericin* allele is not the sole driver of the patterns observed here. While we did not find an effect of the *Diptericin* S69R allele, we cannot rule out a role for *Diptericin* in explaining variation in host susceptibility, although other mechanisms may well also play a role in explaining among species differences in susceptibility. Further study could consider a broader range of species encoding different variants at the S69R site, variation elsewhere in the *Diptericin* gene, or differences in induction of *Diptericin* and other AMPs across species given rapid induction is important for host survival [35, 36, 54].

We found that among species correlations in susceptibility between bacteria all had positive estimates. The strongest significant positive correlation was observed between the two Gram-positive bacteria: *S. aureus* and *E. faecalis*. One explanation for this could be differences in immune defence across the host phylogeny. In *D. melanogaster* Gram-positive bacteria are largely targeted by the Toll pathway [30, 31, 76, 77]. Toll pathway members are highly conserved across arthropods and many Toll pathway genes have large copy number variations across species, which may be linked to their functional diversity [78, 79]. We also found a significant positive correlation across Gram types between *E. faecalis* and *P. rettgeri*. Several studies have demonstrated there is cross-talk between the Toll and IMD pathways, which may play a part in explaining the shared susceptibility [80–82]. However, it is also possible the positive correlations are driven by other factors, for example, the suitability of the host physiological environment for bacterial proliferation.

We found the lowest proportion of variance explained by the phylogeny for *E. faecalis*, for which we also found no correlation between bacterial loads and host mortality. Susceptibility to *E. faecalis* has been shown to involve the action of Bomanin and Baramicin host defence peptides, whose mechanism remains unclear, but could contribute to promoting tolerance to infection rather than pathogen suppression [41, 42, 83]. A previous GWAS study in *D. melanogaster* found variation in susceptibility to *E. faecalis* was due to genes not classically associated with immunity [43], suggesting factors unrelated to bacterial suppression may be important in surviving infection by this pathogen. Further, a study of immune priming showed that flies primed with a low dose of *E. faecalis* were better able to survive subsequent infection by *E. faecalis*, but this was not related to bacterial clearance [84]. *E. faecalis* virulence is known to involve the action of toxins, including Cytolysin and Enterocin V, and induced immune responses including AMPs and iron sequestration have minimal impact on *E. faecalis* virulence [54, 85, 86]. The disconnect between mortality and replication could certainly be explained by toxin-based virulence if low doses of toxins are sufficient to cause mortality as in other Gram-positive insect pathogens [87] and cytolysin, for example, is hemolytic in animal cells in the nanomolar range. Differences in the host response to bacterial toxins such as presence of Toll pathway effector Baramicin A may explain the phylogenetic patterns of susceptibility to *E. faecalis* [42].

We found evidence of microbiome growth in four species (*D. americana*, *S. lebanonensis*, *D. putrida* and *S. pattersoni*) when measuring bacterial loads in homogenized pathogen infected insects. Anecdotally, these species exhibited a relatively high survival to most of the bacteria (Figure 3). The microbiome can play an important role in priming the immune response and defending against bacterial infections [88, 89]. Gut probiotics such as *Lactobacillus* have been demonstrated to reduce fly mortality caused by pathogenic bacteria [90]. It has been reported that there is limited variation in the microbiome across species of Drosophilidae in laboratory culture [91, 92]. As such, a more comprehensive study of how the microbiome varies across these host species, to what extent the microbiome is related to host phylogeny, and whether this relates to differences in susceptibility is warranted.

Although we have studied a relatively large number of host species, only four bacterial taxa were used, limiting our ability to draw broad conclusions on differences between bacterial taxa (for example differences between Gram type). Practical constraints resulted in fewer species being used to estimate the bacterial load data, limiting our ability to assess phylogenetic signal for this trait, or explore the role of tolerance in determining the patterns we observe among host speices. In addition, further work is needed to understand the mechanisms underlying the differences in susceptibility among species, and whether the same or different factors affect susceptibility in different host clades.

In conclusion, we have found that the host phylogeny can explain varying amounts of variation in susceptibility depending on the pathogen. These varying phylogenetic effects (or lack thereof) have important consequences for understanding susceptibility across host species and its impact on pathogen host shifts [26, 61], which is helpful for our understanding of overcoming increasing bacterial infection issues in public health. For some pathogens, certain clades of related hosts will be particularly susceptible, meaning pathogens will be more likely to jump into those species, and they could also act as disease reservoirs [93–96]. However, patterns may be less clear for other pathogens, with relatedness not being a strong predictor of susceptibility across species. As such, caution should be used when estimating the host range of novel pathogens based on susceptibility in closely related host species, without suitable information on patterns of susceptibility across host species for the bacterial taxa in question.

## Supporting information

Supplementary material combined

## Acknowledgements

Bacterial strains were kindly provided by Brian Lazzaro and Ashley Frank. Thanks to Megan Wallace and Michael Jardine for their assistance in the code writing and experimental process for this study. Thanks to editor and two anonymous reviewers for the thoughtful and constructive comments.

**For the purpose of Open Access, the author has applied a CC BY public copyright licence to any Author Accepted Manuscript version arising from this submission.**

## Supplementary materials

### Supplementary methods

**Table S1** Drosophilidae host species, wing length and rearing diet information.

**Table S2** Estimations of the differences between phylogenetic signal of each pathogen pair.

**Table S3** *Diptericin A* sanger sequencing details (primer, information of *DptA* S69R site encoded amino acid, GenBank accession number).

**Table S4** Statistical coefficients for models fitting or not fitting *Diptericin* alleles as fixed effects terms.

**Table S5** Estimates of the mean proportion of variation [Abbot’s corrected mortality, and the day 14 binomial mortality data] explained by the host phylogeny.

**Table S6** Correlations in mortality [Abbot’s corrected mortality, and the day 14 binomial mortality data] between different pathogens.

**Figure S1** Correlations between bacterial load and mortality.

**Figure S2** *Diptericin* allele variation across Drosophilidae species.

**Figure S3** Correlation between mean mortality and Abbot’s corrected mortality.

**All data files and R scripts for analysis used in this study are available in an online repository:** https://doi.org/10.6084/m9.figshare.25648563

## Author Contributions

Conceptualization: HS, BR, BL

Formal Experiments: HS

Data curation: HS, MAH

Formal analysis: HS, RMI, BR, BL

Investigation: HS, MAH, RMI, SKW

Methodology: HS, MAH, BR, BL

Supervision: BR, BL

Visualization: HS, RMI, SKW, BL

Writing- Manuscript draft: HS, MAH, BR, BL

Writing- Reviewing and editing: HS, MAH, BR, BL

