## Supplementary material combined for "Varying phylogenetic signal in susceptibility to four bacterial pathogens across species of Drosophilidae"

**Author information:**

### Supplementary methods

#### Inferring the host phylogeny

Briefly, publicly available sequences of the *28S*, *Adh*, *Amyrel*, *COI*, *COII*, *RpL32*, and *SOD* genes were collected from GenBank (see www.doi.org/10.6084/m9.figshare.6653192 for a full breakdown of genes and accessions by species). Gene sequences were aligned in Geneious v9.1.8 (https://www.geneious.com) using a progressive pairwise global alignment algorithm with free end gaps and a 70% similarity IUB cost matrix. Gap open penalties, gap extension penalties, and refinement iterations were kept as default.

Phylogenetic reconstruction was performed using BEAST version 1.10.4 [1] as the subsequent phylogenetic mixed model (described below) requires a tree with the exact root-tip distances for all taxa. Genes were partitioned into separate ribosomal (*28S*), mitochondrial (*COI*, *COII*), and nuclear (*Adh*, *Amyrel*, *RpL32*, *SOD*) groups. The mitochondrial and nuclear groups were further partitioned into groups for codon position 1+2 and codon position 3, with unlinked substitution rates and base frequencies across codon positions. Each group was fitted to separate relaxed uncorrelated lognormal molecular clock models using random starting trees and four-category gamma-distributed HKY substitution models. The BEAST analysis was run twice, with 1 billion Markov Chain Monte Carlo (MCMC) generations sampled every 100,000 iterations, using a birth-death process tree-shape prior. Model trace files were evaluated for chain convergence, sampling, and autocorrelation using Tracer version 1.7.1 [2]. A maximum clade credibility tree was inferred from the posterior sample with a 10% burn-in. The reconstructed tree was visualised using ggtree version 2.0.4 [3].

#### Sequencing of *Diptericin A*

To determine the *Diptericin A* allele of each species (which has been linked to susceptibility to bacterial pathogens [4]), *Diptericin* genes were detected across species using a recursive tBLASTn approach with a high E-value threshold to compensate for the very short query length (E < 1), followed by manual curation [5]. We then designed primers to amplify *Diptericin A* for 29 species for which sequence data was available (full primer details in Supplementary Table 2). Flies were homogenised in 50 µL DNA extraction buffer (0.5 µL Tris-Cl pH 8.2 (1M), 0.1 µL EDTA (0.5M), 0.25 µL NaCl (5M) and 47.15 µL dd$H_2O$, and 2µL proteinase K at 5mg/µL) then incubated at 37 °C for 30 minutes, followed by 95 °C for 5 minutes. PCRs were then carried out under the following conditions: 95°C for 30 secs, followed by 35 cycles of: 95°C for 20 secs, 54°C for 20 secs, 68°C for 1 minute, followed by a final 5 minutes at 68°C. PCR products were purified using Monarch® PCR&DNA Cleanup Kit, and Sanger sequenced through Eurofins Genomic tube sequencing service. *Diptericin A* S69R site allele analysis was carried out by Geneious prime v10.2.6. All sequences have been uploaded to GenBank (full accession number detailed in Supplementary Table 2).

**Statistical analysis**

Firstly, we included wing size, measured as the length of the IV longitudinal vein from the tip of the proximal segment to the join of the distal segment with vein [6], which provided a proxy for body size [7] and was included in a further model as $wingsize\beta_{2:t}$. This was done to ensure that any phylogenetic signal in body size did not explain the differences in susceptibility between species [8]. Secondly, we ran an additional model which included the effect of polymorphisms at the *Dipterićin A* S69R site ($DptA\beta_{3:t}$) which has previously been implicated in resistance to bacterial pathogens [4, 9, 10]. As variance explained by the phylogeny is calculated after accounting for any fixed effects [11], we examined if variance explained by the phylogeny changed with the inclusion of the *Diptericin* allele (we would expect the phylogenetic variance to decrease if the fixed effects have an effect on mortality and are clustered on the phylogeny).

To validate that the mortality in the control group does not influence our main results, we ran additional models to fit: 1) Abbot's formula corrected mortality; and 2) the mortality on day 14 as the response variable in a binomial model. We used a multivariate model with Abbot's corrected mortality following inoculation with each of the four bacteria as the response variables. Abbot's corrected mortality was calculated as $p_{corr} = \frac{p_{experimental} - p_{control}}{1 - p_{control}}$ [64, 65] where $p_{experimental}$ are vial's mean proportion of dead flies over 14 days, and $p_{control}$ are the mortality in control group of each block. We found a strong positive correlation between raw and Abbot's corrected data (Figure S3). The models took the same form as model (1) and model (2). We used a multi binomial model for the binomial mortality on day 14, the models took the same form as model (1) and model (2). Code for all models is available here (https://doi.org/10.6084/m9.figshare.25648563).

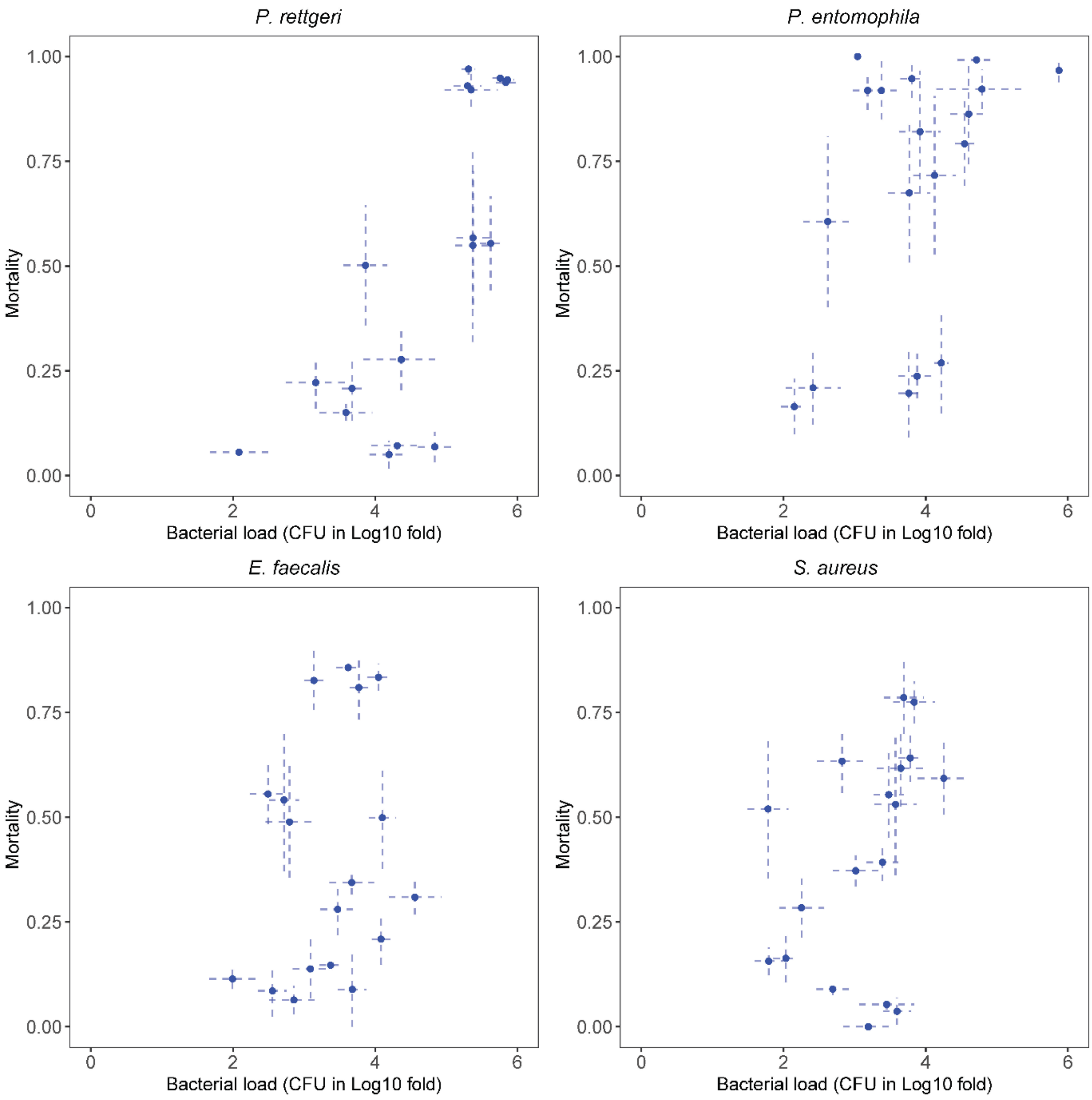


**Figure S1 Correlations between bacterial load and mortality for 18 species of fly.** Each blue dot represents an individual species; the x axis shows the mean bacterial load of individual flies ± SE on a Log 10 scale. The y axis shows the mean mortality ± SE. Correlation coefficients for each pathogen are in Table 4.

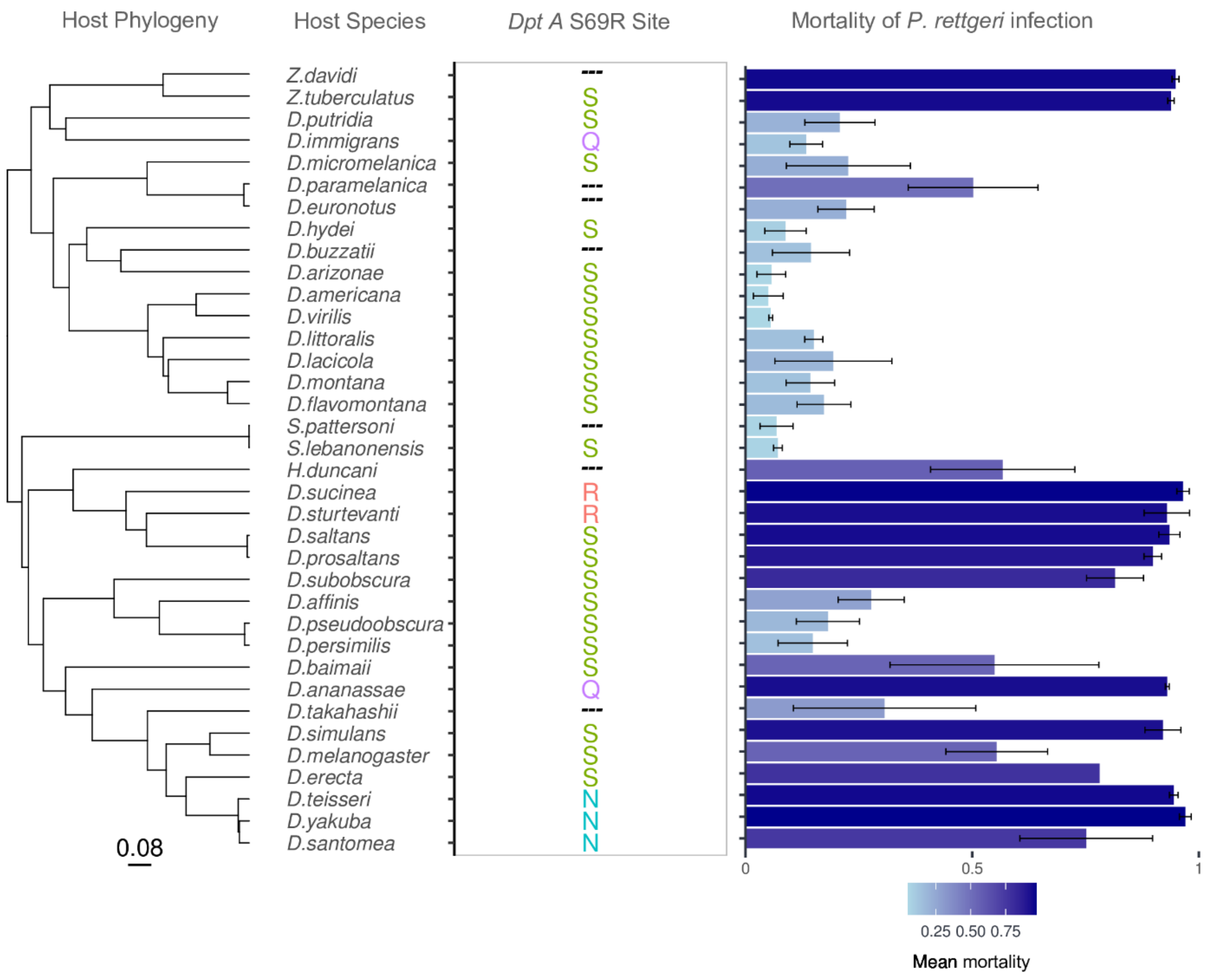


**Figure S2 Diptericin allele variation across Drosophilidae species** Host phylogeny, host species, *Dpt* A allele information and mortality of *P. rettgeri* infection, S= Serine, Q= Glutamine, R= Arginine, N= Asparagine, ---: unknown.

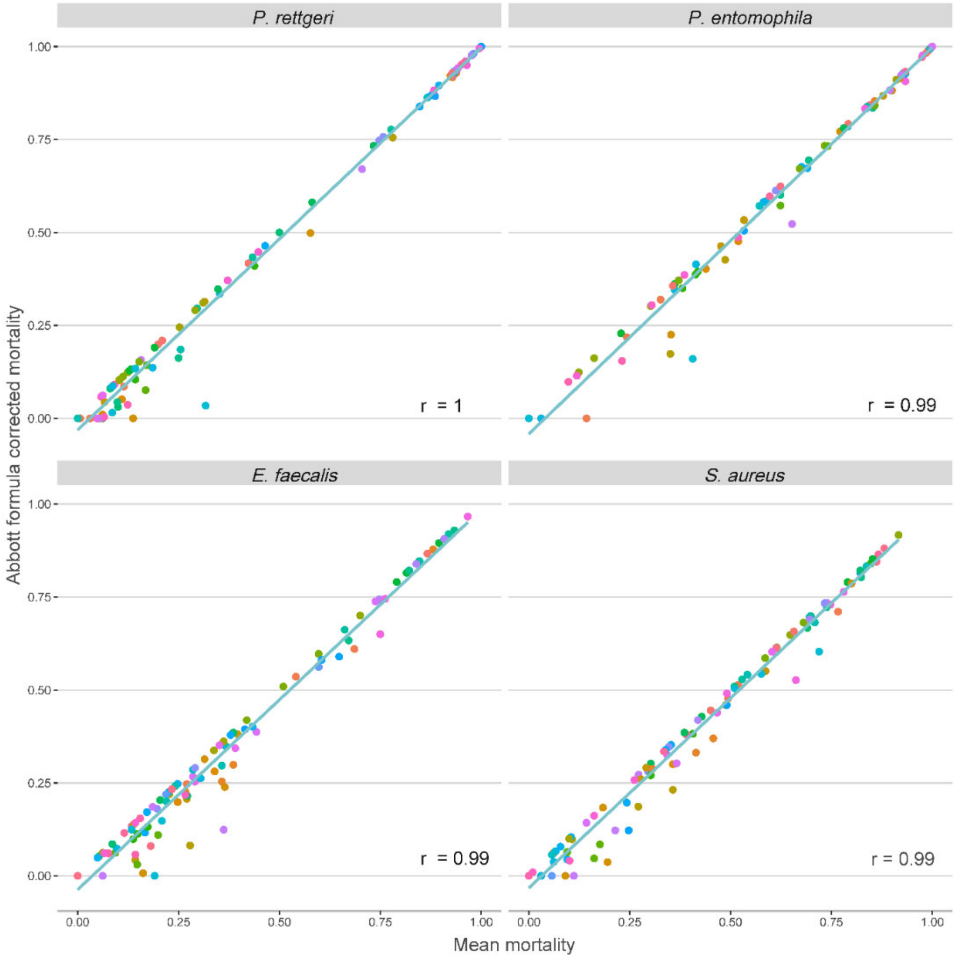


**Figure S3: Correlation between mean mortality and Abbot corrected mortality** Each data point shows the mean mortality (x) and corrected mortality (y) in each pathogen-species-block treatment. r are the Pearson correlation coefficients.

| ID | Animal | Wing length (mm) | Diet | Genus |
|---|---|---|---|---|
| 1 | *D. affinis* | 1.803 | malt | Drosophila |
| 2 | *D. americana* | 2.045 | malt | Drosophila |
| 3 | *D. ananassae* | 1.493 | cornmeal | Drosophila |
| 4 | *D. arizonae* | 1.548 | banana | Drosophila |
| 5 | *D. baimaii* | 1.561 | cornmeal | Drosophila |
| 6 | *D. buzzatii* | 1.902 | malt | Drosophila |
| 7 | *D. erecta* | 1.581 | cornmeal | Drosophila |
| 8 | *D. euronotus* | 2.222 | cornmeal | Drosophila |
| 9 | *D. flavomontana* | 2.192 | malt + yeast | Drosophila |
| 10 | *D. hydei* | 2.182 | cornmeal | Drosophila |
| 11 | *D. immigrans* | 2.153 | malt + yeast | Drosophila |
| 12 | *D. lacicola* | 2.268 | malt | Drosophila |
| 13 | *D. melanogaster* | 1.716 | cornmeal | Drosophila |
| 14 | *D. micromelanica* | 1.895 | cornmeal | Drosophila |
| 15 | *D. montana* | 2.706 | malt + yeast | Drosophila |
| 16 | *D. paramelanica* | 1.946 | cornmeal | Drosophila |
| 17 | *D. persimilis* | 2.013 | malt | Drosophila |
| 18 | *D. prosaltans* | 1.699 | propionic | Drosophila |
| 19 | *D. pseudoobscura* | 1.863 | malt | Drosophila |
| 20 | *D. putrida* | 1.639 | propionic | Drosophila |
| 21 | *D. saltans* | 1.60 | propionic | Drosophila |
| 22 | *D. santomea* | 1.489 | cornmeal | Drosophila |
| 23 | *D. simulans* | 1.484 | cornmeal | Drosophila |
| 24 | *D. sturtevanti* | 1.779 | cornmeal | Drosophila |
| 25 | *D. subobscura* | 2.056 | cornmeal | Drosophila |
| 26 | *D. sucinea* | 1.932 | cornmeal | Drosophila |
| 27 | *D. takahashii* | 1.559 | cornmeal | Drosophila |
| 28 | *D. teisseri* | 1.463 | cornmeal | Drosophila |
| 29 | *D. virilis* | 2.253 | propionic | Drosophila |
| 30 | *D. yakuba* | 1.307 | cornmeal | Drosophila |
| 31 | *H. duncani* | 1.969 | propionic | Hirtodrosophila |
| 32 | *S. lebanonensis* | 2.053 | propionic | *Scaptodrosophila* |
| 33 | *S. pattersoni* | 2.023 | banana | *Scaptodrosophila* |
| 34 | *Z. davidi* | 1.911 | banana | *Zaprionous* |
| 35 | *Z. tuberculatus* | 1.914 | banana | *Zaprionous* |
| 36 | *D. littoralis* | 2.13 | banana | Drosophila |

**Table S1: Drosophilidae host species, wing length and rearing diet information.** Rearing food recipes can be found at https://doi.org/10.6084/m9.figshare.21590724.v1

| Pathogen | Differences in phylogenetic signal |
|---|---|
| ***P. rettgeri - P. entomophila*** | **0.58 (95% CI: 0.17, 0.96)** |
| ***P. rettgeri - E. faecalis*** | **0.80 (95% CI: 0.43, 0.99)** |
| *P. rettgeri - S. aureus* | 0.30 (95% CI: -0.04, 0.69) |
| *S. aureus - P. entomophila* | 0.28 (95% CI: -0.23, 0.82) |
| ***S. aureus - E. faecalis*** | **0.50 (95% CI: 0.02, 0.91)** |
| *P. entomophila - E. faecalis* | 0.21 (95% CI: -0.32, 0.76) |

**Table S2 Estimations of the differences between phylogenetic signal of each pathogen pair** (taken from model (1)). Numbers represent the mean estimates of differences in phylogenetic signal from Model (1) with 95% credible intervals shown in brackets. Significant differences (with 95% CI that do not span zero) are highlighted in bold.

| NO. | Animal | Primer name | Primer sequence | *Dpt A* S 69 R allele information | GenBank accession number |
|---|---|---|---|---|---|
| *1* | *D. affinis* | Daff_DptA- 2 F | GTCTACAATCATCCTCGCTCTGCT | **S** | PP035785 |
| | | Daff_DptA- 291 R | GATAGTTGGTGCCCACGCTG | | |
| *2* | *D. americana* | D.ame-2317F | TTTGTGCATCTCCGACTGCT | S | PP035807 |
| | | D.ame-3299R | TGGAAAATCCGACCTTTTAGGGA | | |
| *3* | *D. ananassae* | D.ana-1226R | AACGCCAAGGACTCTGATGG | Q | PP035789 |
| | | D.ana-444F | TCAGTCGGAAACCTGAAGCC | | |
| *4* | *D. arizonae* | Dari- 146 F | AAAGCGAGTTACCGAAGCCA | S | PP035812 |
| | | Dari- 634 R | TCAGAGGATTTCGTAGTCGAGC | | |
| *5* | *D. baimaii* | D.bai-1056R | CGGGAATCTGAAGCCCAAGT | **S** | PP035804 |
| | | D.bai-393F | GACCACCGATGCGCCTATAA | | |
| *6* | *D. buzzatii* | Dbuz_DptC- 108 R | TCATGTTTAAAAACGGAATCGGTAACT | NA | |
| | | Dbuz_DptC- 112 R | TCATGTTTAAAAACGGAATCGGTAACT | | |
| *7* | *D. erecta* | D.ere-1252R | GCCTGAATGTTCCGGGGTAA | S | PP035799 |
| | | D.ere-341F | AGGTTACGTCGGGGATTCCT | | |
| *8* | *D. euronotus* | Deur_DptC- 345 R | GTAGTTGCCGCCAACACC | NA | |
| | | Deur_DptC- 345 R | GTAGTTGCCGCCAACACC | | |
| *9* | *D. flavomontana* | Dfla_DptC- 39 F | AATCTCGCTGTTCCGGCTAG | S | PP035790 |

| | | | | | |
|---|---|---|---|---|---|
| | | Dfla_DptC- 427 R | AATCGAAATCTGTAGTTGCCACC | | |
| *10* | *D. hydei* | D.hyd-2055F | AAAATTCCCCCAGGCACACA | S | PP035808 |
| | | D.hyd-3669R | TGAAATCCCGCTCAGAGATGA | | |
| *11* | *D. immigrans* | D.immi-2296F | TCTGCTCTGCTCTCGGTACT | Q | PP035805 |
| | | D.immi-3550R | TCAGACTTTGACAGTGCCCA | | |
| *12* | *D. lacicola* | Dlac_DptC_227R | CTAAATCTGTAGTTGCCACCCRCGC | **S** | PP035784 |
| | | Dlac_DptC_4F | CGGAAGCAATTCAATCCGCCAAACG | | |
| *13* | *D. melanogaster* | D.mel_R6-1260R | AGTCTGCCTCAATGTTCCGG | S | PP035800 |
| | | D.mel_R6-418F | AGAGCATCGAAACTGCAGCA | | |
| *14* | *D. micromelanica* | D.mic-2370F | CCCCTCTTCAAACCGATTGC | S | PP035813 |
| | | D.mic-3649R | TGCTCCAAAAGAAGGTCCCA | | |
| *15* | *D. montana* | D.mon-2328F | TCGGCTGGTGCACCTTTTAT | S | PP035814 |
| | | D.mon-3597R | CAGACCCAGCCCAGGTTAG | | |
| *16* | *D. paramelanica* | Dpar_Dpt_-1F | CAATATGGTTCGACAGCGA | NA | |
| | | Dpar_Dpt_-130R | TCAGCTTTGTTCAAGTCACA | | |
| *17* | *D. persimilis* | Dper-1263R | CCAAGAAGGGTCTCGGATCG | **S** | PP035791 |
| | | Dper-389F | AGGGACCACAATCCGGACTA | | |
| *18* | *D. prosaltans* | Dpro_DptA_595R | GTCATAAATGTTCCGGGTTAAAGAAACT | **S** | PP035792 |
| | | Dpro_DptA_139F | AAAAGGGTCCACAATCCAATTGG | | |

| | | | | | |
|---|---|---|---|---|---|
| *19* | *D. pseudoobscura* | Dpse-1277R | TGGAAAGTCCTCCCCCAAGA | S | PP035793 |
| | | Dpse-389F | AGGGACCACAATCCGGACTA | | |
| *20* | *D. putridia* | Dput_DptC- 701 R | AAATCTGAATCTGTAGATACCGCC | S | PP035787 |
| | | Dput_DptC- 2 F | ATCCTTGCTTTGGTCTGCTTC | | |
| *21* | *D. saltans* | Dsal-1392R | TCAAGCAACCATAAAGCACAGA | S | PP035794 |
| | | Dsal-200F | GGCAAATGTGGTGCATTGGA | | |
| *22* | *D. santomea* | Dsan-1252R | GCCTGAATGTTCCGGGGTAA | N | PP035801 |
| | | Dsan-426F | GCAACTGCAGCAGAGCTATC | | |
| *23* | *D. simulans* | Dsim_w501-1252R | GCCTCAATGTTCCGGGGTAA | S | PP035798 |
| | | Dsim_w501-213F | GCCGTGGGTGTACTTTCTGA | | |
| *24* | *D. sturtevanti* | Dstu-1394R | ACTCAAGCAACCATAAAGCACA | R | PP035795 |
| | | Dstu-365F | AGGGGGACTTACTTTTGCACC | | |
| *25* | *D. subobscura* | Dsubo-1332R | ATCATCCAAAGCCCCGTCTC | S | PP035797 |
| | | Dsubo-431F | TGCAGCCATGGCATCAGTTA | | |
| *26* | *D. sucinea* | Dsuc-1242R | TCCGGGGTAAATGAACGTGT | R | PP035796 |
| | | Dsuc-206F | TGTGGTTCATGTGGGGGATT | | |
| *27* | *D. takahashii* | Dtak-1240R | GTGACCGACCGACGATCTTT | **NA** | |
| | | Dtak-114F | CGGGGTAAACAAACAACGCC | | |
| *28* | *D. teisseri* | Dtei-1252R | GCCAGAATGTTCCGGGGTAA | N | PP035802 |

| | | | | | |
|---|---|---|---|---|---|
| | | Dtei-431F | TGCAGCAGAGCTATCAGTCG | | |
| *29* | *D. virilis* | Dvir_RS2-2332F | CTGCTGCGTGGTCTAGCTAT | S | PP035811 |
| | | Dvir_RS2-3224R | ACGAATCCCCAAACTTGTGT | | |
| *30* | *D. yakuba* | Dyak-1309R | CAGCCTCCGTCTCAGTCTCT | N | PP035803 |
| | | Dyak-407F | CGGCCTATAAAAGCGCATCG | | |
| *31* | *H. duncani* | | Unknown | | |
| *32* | *S. lebanonensis* | Sleb_DptA_RS2_1247R | CCAGTTCCGGGTTTGGTGAA | **S** | PP035786 |
| | | Sleb_DptA_RS2_259F | CTGCTGTGCTGGCCTATCC | | |
| *33* | *S. pattersoni* | Sleb_DptA_RS2_1247R | CCAGTTCCGGGTTTGGTGAA | NA | |
| | | Sleb_DptA_RS2_259F | CTGCTGTGCTGGCCTATCC | | |
| *34* | *Z. davidi* | Zdav-2174F | ATAAGGCTGGACAGAGGGGT | NA | |
| | | Zdav-3660R | ACTCAAGGATCACTCTCAAAGA | | |
| *35* | *Z. tuberculatus* | Ztub_DptC_515R | TTCACAAATAGTACATCAAAATCTAAAGCG | **S** | PP035788 |
| | | Ztub_DptC_171F | CTGCAGCATCAGTTAATCAGTCAG | | |
| *36* | *D. littoralis* | D.lit-2314F | GGTTCTGTGCATCTCCGACT | S | PP035809 |
| | | Dlit-3362R | TTGGGCTCACGTGTTGCATA | | |

**Table S3 *Diptericin A* sanger sequencing details (primer sequence, *Dpt A* S69R encoded amino acid, GenBank accession number)**

| Pathogen | Change in mortality in presence of susceptible *Dpt A* allele | Change in phylogenetic signal |
|---|---|---|
| *P. rettgeri* | 0.21 (95% CI's -0.12, 0.52) | 0.00 (95% CI's -0.22, 0.18) |
| *P. entomophila* | 0.29 (95% CI's -0.02, 0.54) | -0.11 (95% CI's -0.64, 0.50) |
| *E. faecalis* | 0.24 (95% CI's -0.03, 0.50) | 0.01 (95% CI's -0.51, 0.63) |
| *S. aureus* | 0.07 (95% CI's -0.20, 0.33) | 0.01 (95% CI's -0.57, 0.61) |

**Table S4 Effect of *Diptericin A* alleles on susceptibility.** The difference in mortality shows the effect of *Dpt* A allele (resistant or susceptible) when included in the model as a fixed effect. The change in phylogenetic signal shows the difference in the proportion of variation in mortality explained by the phylogeny between models not including vs including Diptericin alleles as fixed effects. If the *Dpt* A allele status was explaining the observed patterns we would expect to see a significant decrease in the proportion of variation explained by the phylogeny. Numbers represent the mean estimates with 95% credible intervals shown in brackets.

| Pathogen | Abbot's corrected mortality | Day14 binomial data |
|---|---|---|
| ***P. rettgeri*** | **0.93 (95% CI: 0.80, 1.00)** | **0.93 (95% CI: 0.79,1.00)** |
| ***P. entomophila*** | **0.29 (95% CI: 0.00, 0.64)** | **0.41 (95% CI: 0.00, 0.79)** |
| ***E. faecalis*** | **0.14 (95% CI: 0.00, 0.52)** | **0.18 (95% CI: 0.00, 0.60)** |
| ***S. aureus*** | **0.65 (95% CI: 0.26, 0.97)** | **0.74 (95% CI: 0.45, 0.97)** |

**Table S5 Estimates of the mean proportion of variation [Abbot's corrected mortality, and the day 14 binomial mortality data] explained by the host phylogeny (taken from model (1)).**

| Bacteria | Abbot's corrected mortality | Day 14 binomial data |
|---|---|---|
| ***S. aureus - E. faecalis*** | **0.67 (95% CI: 0.39, 0.90)** | **0.69 (95% CI: 0.46, 0.89)** |
| *S. aureus - P. rettgeri* | 0.29 (95% CI: -0.01, 0.66) | **0.50 (95% CI: 0.16, 0.76)** |
| *S. aureus - P. entomophila* | 0.40(95% CI: -0.03, 0.79) | 0.16 (95% CI: -0.31, 0.57) |
| ***E. faecalis - P. rettgeri*** | **0.42 (95% CI: 0.06, 0.75)** | **0.47 (95% CI: 0.16, 0.76)** |
| *E. faecalis - P. entomophila* | 0.39(95% CI: -0.02, 0.82) | 0.09 (95% CI: -0.33, 0.51) |
| ***P. rettgeri - P. entomophila*** | **0.42 (95% CI: 0.05, 0.78)** | 0.08 (95% CI: -0.33, 0.48) |

**Table S6 Correlations in mortality [Abbot's corrected mortality, and the day 14 binomial mortality data] between different pathogens.** Numbers represent the mean estimates of correlation coefficients from Model (2) with 95% credible intervals shown in brackets. Significant correlation coefficients (with 95% CI that do not span zero) are highlighted in bold.
